# Multi-locus metabarcoding and intensive sampling reveal extraordinary diversity carried in the ballast water of a single vessel

**DOI:** 10.64898/2026.05.07.723533

**Authors:** Sarah Brown, Katharine J. Carney, Katrina M. Pagenkopp Lohan, Kimberly K. Holzer, Erik M. Pilgrim, Gregory M. Ruiz, John A. Darling

## Abstract

Understanding risks of biological invasions associated with ballast water (BW) requires full understanding of the biodiversity transported in ballast tanks. Here we characterize the remarkable level of diversity that can be carried in the BW of a single vessel. To maximize our ability to capture BW diversity we: 1) utilized DNA-based methods to describe biodiversity, including both native and non-native taxa; 2) exploited multiple primer sets targeting multiple genomic loci with different expectations for taxonomic coverage; 3) assessed multiple tanks on a single vessel to capture different communities present in different tanks; and 4) sampled those tanks with far higher-than-usual replication both to improve representation of the diversity present and to enable statistical estimation of total richness. Using this approach, we found extraordinarily high diversity associated with a single vessel. Across all loci, we estimate a total of 272 taxa that can be assigned species names; looking more broadly at unnamed molecular operational taxonomic units, our estimates are between 425 and 742 individual taxa, depending on the locus. We confirm that only a fraction of this diversity would be captured with typical sampling efforts. We found that different loci capture different snapshots of biodiversity and that our ability to detect taxa of interest (e.g., non-native species) depends on sampling effort and genomic locus. Our results expand upon previous studies describing highly diverse BW communities and add to a growing literature that demonstrates the value of molecular methods for characterizing those communities and assessing the associated risk of non-native species introduction.

## 1. Introduction

How much biodiversity is there in a ballast water tank? This question has been of interest for as long as scientists have contemplated the risks of BW-borne biological invasions. In 1993, based on one of the first surveys of BW biodiversity, Carlton & Geller suggested that on any given day “several thousand” species may be moving around the world’s oceans in ballast holds. This they derived from an approximation of “20 to 30 taxa per vessel” [1] while recognizing that the number was likely an underestimate given limitations on their ability to identify larval stages and the likelihood that many small-sized taxa evaded capture, among other things.

Such early observations led to a decades-long conversation about the role BW might play in driving anthropogenic translocations of aquatic species and associated biological invasions with potentially catastrophic ecological and economic costs [2-8]. Recognition of these risks ultimately drove establishment of multiple regulatory frameworks designed to reduce the negative impacts of BW-borne introductions. These include the International Maritime Organization’s (IMO) 2004 International Convention for the Control and Management of Ship’s Ballast Water and Sediments [9], which entered into force in September 2017 [10], as well as regulations adopted by individual nations, including those established by the U. S. Coast Guard [11] and U. S. Environmental Protection Agency [12].

Improved characterization of BW-borne biodiversity facilitates assessment and mitigation of risks posed by BW-driven species introductions. Numerous early efforts assessed portions of the diversity in BW tanks, in various contexts and focusing on multiple taxonomic groups [13-15] and large assemblages of vessels [16, 17]. However, such studies were constrained by the same caveats noted by Carlton & Geller (1993): namely, morphological analysis is restricted to those taxa for which good morphological keys (and good taxonomists) exist, and to those individuals that exhibit the characteristics referenced in those keys [18]. The advent of DNA metabarcoding has enabled more expansive estimates of the biodiversity present in BW, in part by capturing some of the diversity hidden to morphological taxonomy [19, 20]. Metabarcoding studies have now dramatically increased estimates of the number of taxa present in BW samples, with sampling efforts across multiple vessels suggesting that hundreds [21, 22] of operational taxonomic units (OTUs, broadly understood to be roughly equivalent to species-level diversity) are regularly transferred among ports in BW [23].

While the application of metabarcoding methods has clearly led to more generous estimates of the total diversity present in BW, the approach is burdened with its own caveats. Among the most obvious of these are that OTUs may not be equivalent to species [28-30], and that gaps in the reference databases used to assign taxonomic names to sequences result in incomplete or uncertain identifications for much of the data generated in metabarcoding studies [31]. As an example, a recent study looking at dinoflagellate diversity in BW sediments uncovered an astonishing 2410 OTUs across 49 samples; however, after bioinformatic analysis only 33 species names could be confidently assigned [32]. Additionally, the choice of amplicon marker region and primers can result in substantially different results in detected biodiversity. The two most commonly used “universal” eukaryotic marker regions, the 18S ribosomal RNA (rRNA) gene and the mitochondrial cytochrome C oxidase subunit I (COI) gene region, are both effective at detecting a wide range of eukaryotic taxa and thus have become standard marker regions used to detect eukaryotic biodiversity [33, 34]. However, each region has limitations in its detection capabilities – for example, the 18S region may provide less resolution down to the species level for some taxa [35], while targeting the COI region can result in gaps in taxonomic coverage due to the relatively fast mutation rates of the COI region [36]. Incorporating the use of multiple metabarcodes into a biodiversity study can provide an effective solution to the biases introduced as a result of region or primer choice by enhancing taxonomic coverage and resolution, ultimately enabling greater detections of overall biodiversity [35, 37, 38].

While studies examining biodiversity have begun to utilize multi-locus metabarcoding approaches for more comprehensive estimates of biodiversity [37-39], few examples of such multi-locus metabarcoding methods being utilized in BW biodiversity studies exist [24, 25, 40, 41]. Despite a limited number of samples, existing BW biodiversity studies that utilize multi-locus metabarcoding have so far revealed the presence of substantial biodiversity across phylogenetic groups. For instance, Ardura et al. explored changes in BW diversity over the course of multiple 28-day voyages using COI and 18S metabarcoding and found 165 OTUs assigned to 137 genera on one trip and 631 OTUs assigned to 337 genera on a second [24]. Similarly, an examination of three BW samples taken from a single ship at multiple time-points during a 3-week cruise identified 29 genera using the COI marker and 136 genera using the chloroplast RuBisCo large subunit (rbcl) across multiple metazoan and algal groups [25]. Notably, when multi-locus metabarcoding of the 18S and 12S genomic regions was used to investigate the presence of species across multiple taxonomic groups in BW, 76 phytoplankton species, 70 invertebrate species, and 100 fish species were successfully identified [40].

Although these studies have provided an improved sense of the potential scale of biodiversity transported in BW, to our knowledge no existing study has attempted explicitly to provide an estimate of the total eukaryotic diversity present in a single vessel and point in time. In part, this is certainly due to the difficulties associated with sampling BW tanks. Obtaining samples can be logistically challenging, and time and resource constraints typically limit studies to one or a few samples per vessel except in rare cases (e.g. see [14, 15, 24, 42]). Given the known patchiness of BW biodiversity [43, 44], standard sampling efforts are therefore highly unlikely to provide good estimates of total diversity in BW tanks. In fact, the challenge of obtaining representative samples has led to establishment of protocols specifically designed to overcome these limitations and provide more accurate assessments of organismal densities to test compliance of BW management systems (BWMS) with numerical discharge standards [44, 45]. However, these sampling approaches aim principally to assess abundance (concentrations) of live organisms in compliance contexts and have not generally been geared toward estimation of total biodiversity.

Here we describe the results of an intensive sampling effort of the BW of a single ship entering the port of Valdez, Alaska, USA. To achieve more representative sampling of BW and, more importantly, to enable statistical estimations of total biodiversity, we collected 10 plankton net tow samples in each of two BW tanks, for a total of 20 samples. We maximized the taxonomic breadth of our assessment and our ability to detect rare or typically unacknowledged (e.g., single-celled eukaryotic) taxa given limited resources by adopting a multi-locus metabarcoding approach targeting multiple regions of the COI and 18S gene regions. Specifically, we chose primers that amplify a wide variety of taxa (Table 1) in order to enhance richness detection [46] and utilized different primer sets for 18S to amplify the same genomic region in order to account for any gaps in taxonomic coverage introduced by different primers. Our primary objective was to capture as many eukaryotic species as possible present in the BW of a single ship and to assess the value of molecular methods for describing this diversity. Given the obvious importance of introduced taxa in the context of BW transport, we also examine the recovered biodiversity for the presence of non-native species and explore differences in the detection of both native and non-native diversity across different genomic markers. We further discuss the benefits and limitations of these molecular methods, with respect both to the overall estimation of biological diversity and to the detection and identification of non-native taxa in BW.

**Table 1.**
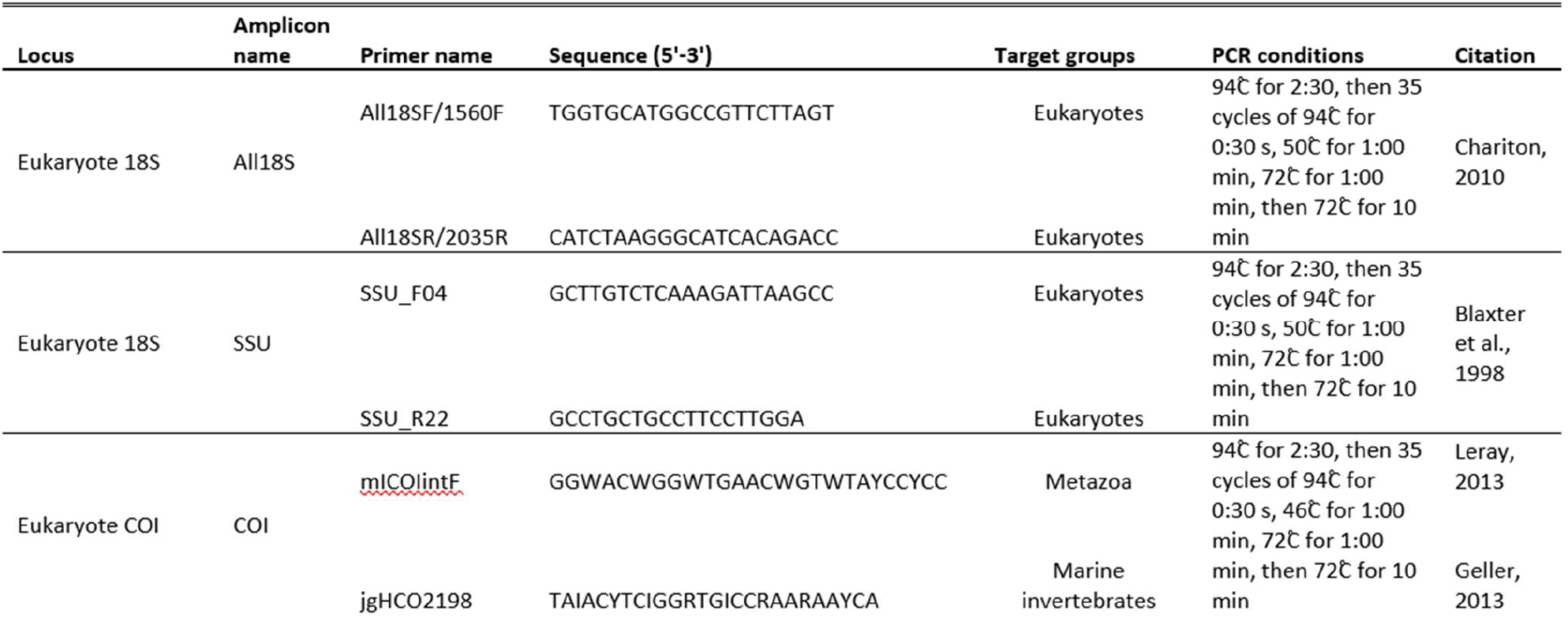
Primers, PCR conditions, and references for each 18S and COI amplicon targeted in this study.

## 2. Materials and Methods

### 2.1 BW sampling

BW was sampled as described previously [26] from a crude oil tanker arriving at the Valdez Marine Terminal in Alaska. Two ballast tanks were accessed and 10 samples per tank were taken using a plankton net with 80 μm mesh. The cod end of the net was lowered to the lowest point accessible in the tank and towed vertically to the surface at a consistent speed. Biomass from the net and cod end was rinsed into a 125 mL Nalgene bottle using filtered tank water and transported on ice back to the laboratory where samples were re-filtered to remove water and preserved in 95% ethanol in 50 mL Falcon tubes. Prior to DNA extraction, preserved samples were further concentrated onto a 20 μm mesh filter and rinsed with ethanol.

### 2.2. DNA extraction, library preparation, and sequencing

Phenol-chloroform DNA extractions were performed on filters as described previously [47]. We amplified two regions of the 18S ribosomal RNA (rRNA) and one region of the cytochrome C oxidase subunit I (COI) genes using three sets of primers (Table 1). Amplification conditions for each set of primers are provided in Table 1. Dual indexing and 2×300 paired-end Illumina MiSeq sequencing were conducted according to standard protocols described in further detail in [47]. Sequences have been deposited in the National Center for Biotechnology Information (NCBI) Sequence Read Archive (SRA) under accession number PRJNA1271602.

### 2.3. Bioinformatics

To process COI and 18S rRNA sequence data, demultiplexed paired-end FASTQ files were imported into QIIME2 (version 2023.7) [48] and merged, trimmed, denoised, and dereplicated using DADA2 [49]. To assign taxonomy to COI sequences, a custom COI reference sequence database was created using mkCOInr (version 0.3.0) and the COInr database (version 2024_05_06) [50]. Briefly, the COInr database, which contains COI sequences from both NCBI and BOLD, was filtered to create a custom database which contained only sequences amplified with the COI primers after non-Eukaryote sequences were removed from the database. Only sequences that covered a minimum of 80% of this region were included in the final database. Additionally, taxa from the class Insecta were subsequently removed from the custom database to reduce database size and to avoid false taxonomic assignments, as insect diversity is generally low in marine environments and not the focus of this study [51]. The resulting COInr database for the COI amplicon contained 910,547 reference sequences, respectively, and was formatted for use in QIIME2 (version 2023.7).

For 18S taxonomy assignment, primer-specific reference sequence databases were created from the Silva 138 SSURef NR99 full-length sequence database [52] using the ’extract-seq-segments’ command in RESCRIPt [53] to perform an iterative reference sequence extraction process. The final reference sequence databases for sequences amplified using the all18S and SSU primers contained 39,874 and 40,948 sequences, respectively.

Three Naïve Bayes classifiers were trained on these custom databases using QIIME2’s q2-feature-classifier plugin [54], and taxonomy was assigned to query sequences using default parameters. All code written for bioinformatic analyses and custom reference database curation is available on GitHub (https://github.com/sbrown2648/BW-multi-locus-metabarcoding).

### 2.4. Data analysis

For statistical analyses, QIIME2 artifacts were imported to Rstudio (version 2024.9.0.375) [55] using the qiime2R package (version 0.99.6) [56]. Prior to additional analysis, samples with fewer than 10,000 reads were removed from the dataset using the phyloseq package (version 1.48.0) [57]. The remaining sample sizes per marker were 19 for all18S samples, 20 for SSU, and 15 for COI. Samples were rarified to 14,156 reads (the lowest number of reads observed in any included sample) to eliminate the impact of sequencing depth on comparisons between loci.

We adopted multiple approaches to estimating overall observed taxonomic richness. To estimate taxonomically un-assigned molecular biodiversity, we clustered ASVs into operational taxonomic units (OTUs) at a 97% similarity cutoff using the R package speedyseq (version 0.5.3.9021) [58] in order to approximate species-level richness as commonly reported in existing literature on BW diversity. This approach has disadvantages over direct analysis of ASVs in terms of overall diversity assessment; for instance, it precludes identification of intraspecific diversity. However, it usefully allows comparison with other studies that report species-level diversity, including those based on morphological analyses, because it enables assessment of the total OTU richness. Since this includes those OTUs that cannot be assigned taxonomic names, it provides the most liberal assessment of such diversity that is available to us. However, the approach does not allow for joint evaluation of richness across all loci combined. To accomplish that, we also assigned taxonomic names to unclustered ASVs at the highest resolution possible. ASVs were then agglomerated by taxonomy, allowing comparison of composition across loci at multiple levels of taxonomic resolution. This approach enabled us to pool across loci to estimate total richness of named species, with the obvious disadvantage of discarding taxa that could not be assigned at the species level.

Rarefaction and extrapolation curves were created using iNEXT (version 3.0.1) [59]. We pooled all samples across both tanks for rarefaction analyses, as we observed no significant differences in community composition between tanks (see Results). To estimate sample coverage, the incidence frequency function in the iNEXT package was used to calculate the percent coverage per number of samples (maximum of 5 samples) and the number of samples needed to reach 50%, 90%, 95%, 99%, and 100% of the total estimated biodiversity. To examine differences in alpha diversity between groups, values for richness (observed) and Shannon Diversity were calculated in phyloseq and evaluated using pairwise Wilcoxon Rank Sum tests with the Benjamini-Hochberg adjusted p-value method (stats package, version 4.4.0) [60]. Diversity accumulation curves were generated independently for all loci using OTUs clustered at 97% similarity as described above. We also generated additional curves using iNEXT based on the reduced dataset including only ASVs taxonomically assigned names at the species level.

Prior to analysis of community composition, phylum names were compared across markers and updated to reflect the taxonomic nomenclature used in the World Register of Marine Species database. Visualizations of differences in community composition between markers were conducted with ASVs assigned at both the phylum and species level using the metaMDS command in the vegan package (version 2.6-6.1) to produce non-metric multidimensional scaling (NMDS) plots based on the Bray-Curtis ordination [61]. To determine whether variations in community composition were significant, PERMANO-Vas (Permutational Multivariate Analysis of Variance) were computed using the adonis2 function in vegan. Further visualization of phylum and species level community composition was conducted using multi-set intersection analyses in the SuperExactTest package (version 1.1.0) [62]. To identify non-native species, a list of all species with a history of introduction to North America were compiled from the National Exotic Marine and Estuarine Species Information System (NEMESIS) database [63] (accessed October, 2024) and cross-referenced with the species level identifications of the three markers.

The files, metadata, and code used in this analysis are available on GitHub (https://github.com/sbrown2648/BW-multi-locus-metabarcoding).

## 3. Results

Overall patterns in diversity and sample coverage estimates did not differ notably between tanks (Fig. S1); neither did Shannon diversity or richness, which did not exhibit significant differences between tanks (Wilcoxon Rank Sum Test, p > 0.05) (Fig. S2, Table S1). Tank-specific differences in OTU level community composition were apparent only when samples were sequenced with the COI primers, where differences were small but statistically significant (Fig. S1; PERMANOVA, R^2^ = 0.1618, *p* = 0.02). These results suggested that samples could be pooled across tanks for overall diversity analyses.

In total, we obtained 184 identifiable species belonging to 28 eukaryotic phyla across all 3 primer sets; however, not all ASVs could be identified to the species level, so overall estimated diversity was far higher. Estimated total diversity within the BW tanks and the number of samples required to capture that biodiversity varied across markers (Fig. 1, Tables 2, 3). Total estimated OTU richness for the COI primer set was 742 (95% CI from 643 to 841); for all18S it was 544 (95% CI from 498 to 589); and for SSU it was 425 (95% CI from 344 to 507; all estimates are rounded to the nearest whole number). Even the intensive sampling approach adopted here was insufficient to achieve 95% sample coverage for any of the markers (Table 2). Rarefaction analysis indicated that the intensive sampling conducted in this study was only sufficient to capture most of the total estimated diversity for each primer set (88% for all18S with 19 samples, 87% for COI with 15 samples, and 90% for SSU with 20 samples), as curves approached plateau with the number of samples taken (solid line in Fig. 1b). The diversity captured by a single BW sample–a standard sampling effort given typical resource limitations–was roughly half of the expected biodiversity for COI and SSU, but only 10% for all18S (Table 3). Even 5 samples were insufficient to capture 90% of the expected diversity for any of the markers. When restricting analyses to only those ASVs that could be assigned species names, we still identified considerable diversity across all three primer sets (Fig. 1C), with species-level diversity being substantially higher for the COI marker. This analysis allowed us to combine data across all 3 loci; doing so we conservatively estimated total taxonomically identified species-level diversity at 272 species (95% confidence interval 223 to 321; Fig. 1E).

**Figure 1.**
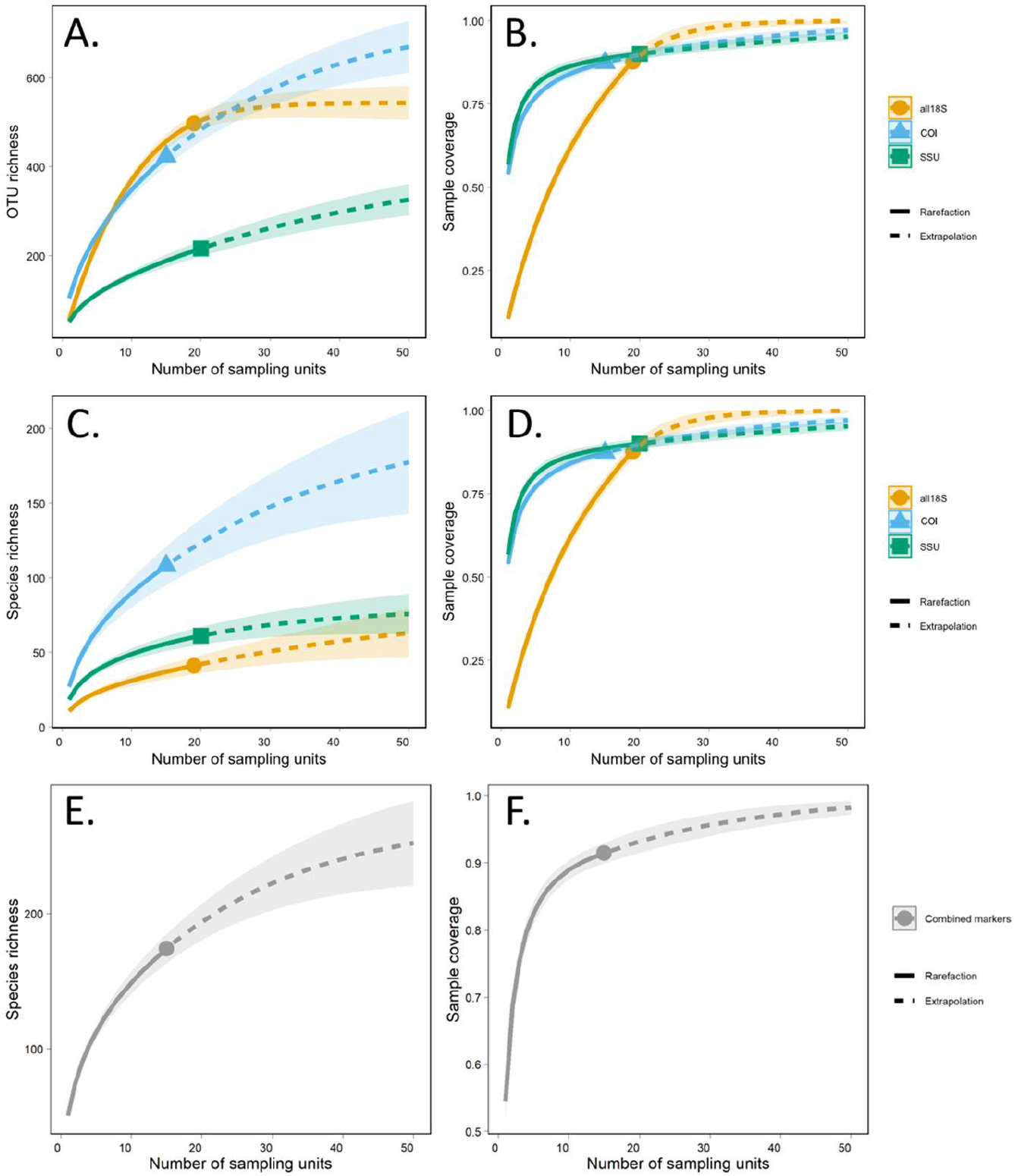
Results of iNEXT analysis based on data rarefied at 14,156 reads. A. Diversity accumulation curve, with OTU diversity (clustered at 97%) on the y axis and number of sampling units on the x axis. B. Sample coverage, with the fraction of total estimated species diversity on the y axis and number of sampling units on the x axis. Data for different loci are plotted separately, as shown in the legend. Solid lines indicate interpolation based on observed data, dashed lines represent extrapolated estimates, shaded areas show 95% confidence intervals. C and D. As in A and B, but data include only ASVs identified taxonomically at the species level. E and F. Accumulation and coverage curves using only ASVs identified taxonomically at the species level, with data from 3 loci combined.

**Table 2.** Number of samples required to achieve 50%, 90%, 95%, and 99% coverage of total estimated species-level diversity for each locus. Asterisks indicate estimates with high uncertainty (estimated number of samples more than twice the observed number). Number of samples rounded to nearest whole number.

| Locus | Percent coverage |  |  |  |
| --- | --- | --- | --- | --- |
|  | 50 | 90 | 95 | 99 |
| **all18S** | 7 | 20 | 25 | 35 |
| **SSU** | 1 | 18 | 48* | 113* |
| **COI** | 1 | 20 | 37* | 75* |

**Table 3.** Percent of total estimated species-level diversity captured by sample sizes n = 1 through n = 5. Percentages are rounded to nearest whole number.

| Locus | Number of sampling units |  |  |  |  |
| --- | --- | --- | --- | --- | --- |
|  | 1 | 2 | 3 | 4 | 5 |
| **all18S** | 10 | 19 | 26 | 33 | 39 |
| **SSU** | 57 | 69 | 75 | 78 | 81 |
| **COI** | 54 | 65 | 71 | 74 | 77 |

We observed significant marker-based differences in diversity measures. OTU richness was significantly higher in the COI dataset (median = 98) than for both 18S amplicons (Pairwise Wilcoxon Rank Sum Test, p < 0.05), and Shannon diversity exhibited significant differences between all amplicons (Fig. S3, Table S1). An NMDS analysis of community composition at the phylum level revealed some overlap in clustering of the three markers, though differences at this taxonomic level were significant (Fig. 2a; PERMANOVA, R^2^ = 0.6278, *p* = 0.001). The recovered taxa consisted of 18 phyla in the all18S dataset, 23 phyla in the SSU dataset, and 21 phyla in the COI dataset (Fig. S4, Fig. 2b). Of these, fourteen phyla were observed across all markers, including Annelida, Arthropoda, Bryozoa, Chaetognatha, Chordata, Cnidaria, Echinodermata, Haptophyta, Heterokontophyta, Mollusca, Myzozoa, Nematoda, Nemertea, and Platyhelminthes (Fig. 2b, Fig. S4).

**Figure 2.**
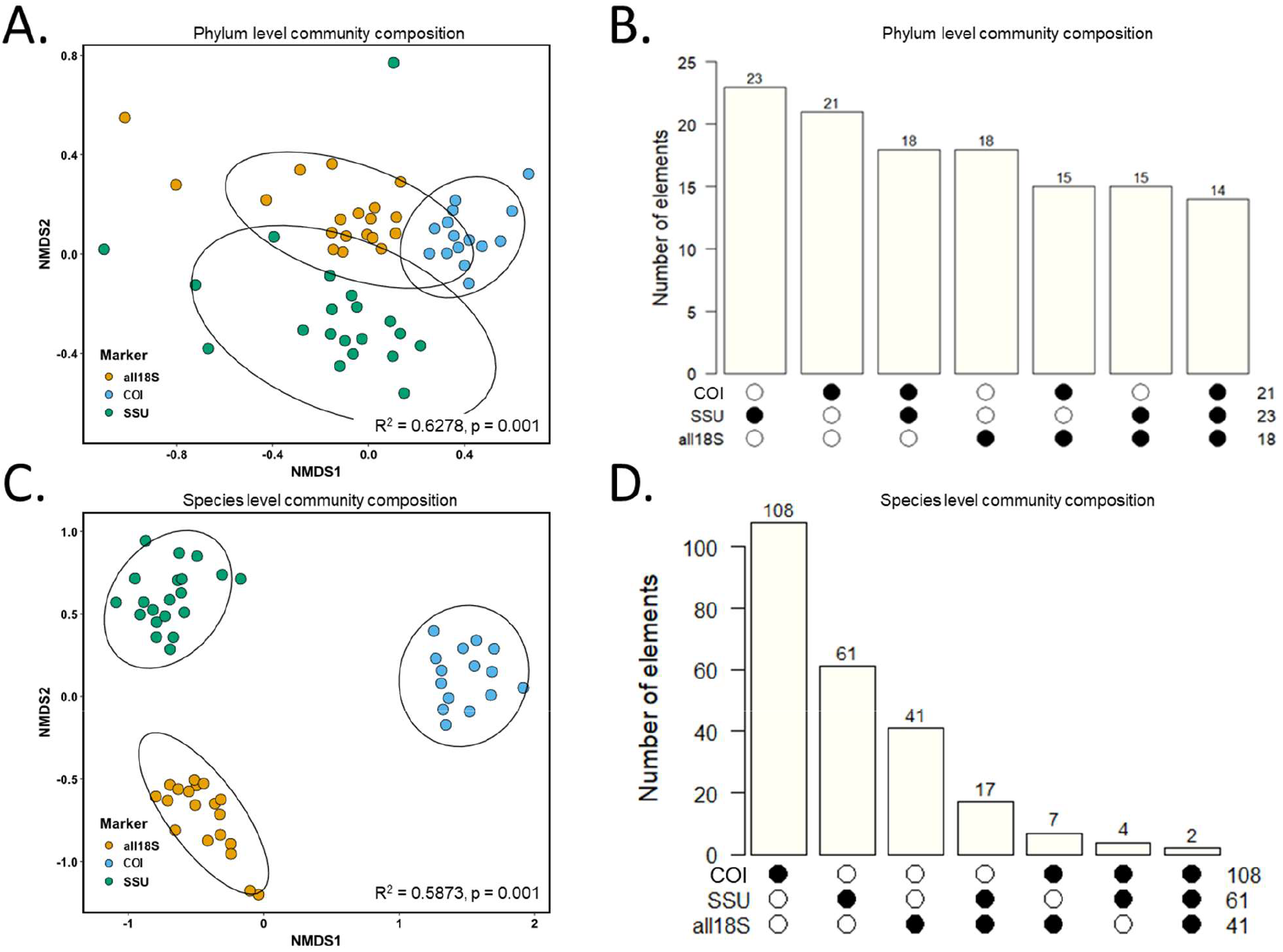
Analysis of overlap of taxonomic coverage by different loci. Non-metric multidimensional scaling (NMDS) plot based on the Bray-Curtis ordination of community composition assessed at the phylum level (A), and the species level (C) based on rarefied data. Ellipses indicate 95% confidence intervals. Intersection plots showing overlap of the phyla (B) and species (D) captured by the three genomic loci, with the x-axis representing the markers for which the phyla or species are present. Note that the total number of taxa detected for each individual loci include phyla or species that are also detected by other markers. Taxa unassigned at the phylum (A, B) or species (C, D) levels were removed prior to analysis.

As at the phylum level, community composition varied significantly between amplicons at the species level (Fig. 2c; PERMANOVA, R^2^ = 0.5873, *p* = 0.001). However, species-level analyses revealed greater differentiation between the three markers (Fig. 2c and 2d). The number of taxa resolved to the species level varied substantially across the three markers (Fig. 2d and Table 4). An analysis of species level taxonomic assignments revealed that the COI marker had the highest number of species level identifications, while the all18S marker had the fewest (Table 4).

**Table 4.**
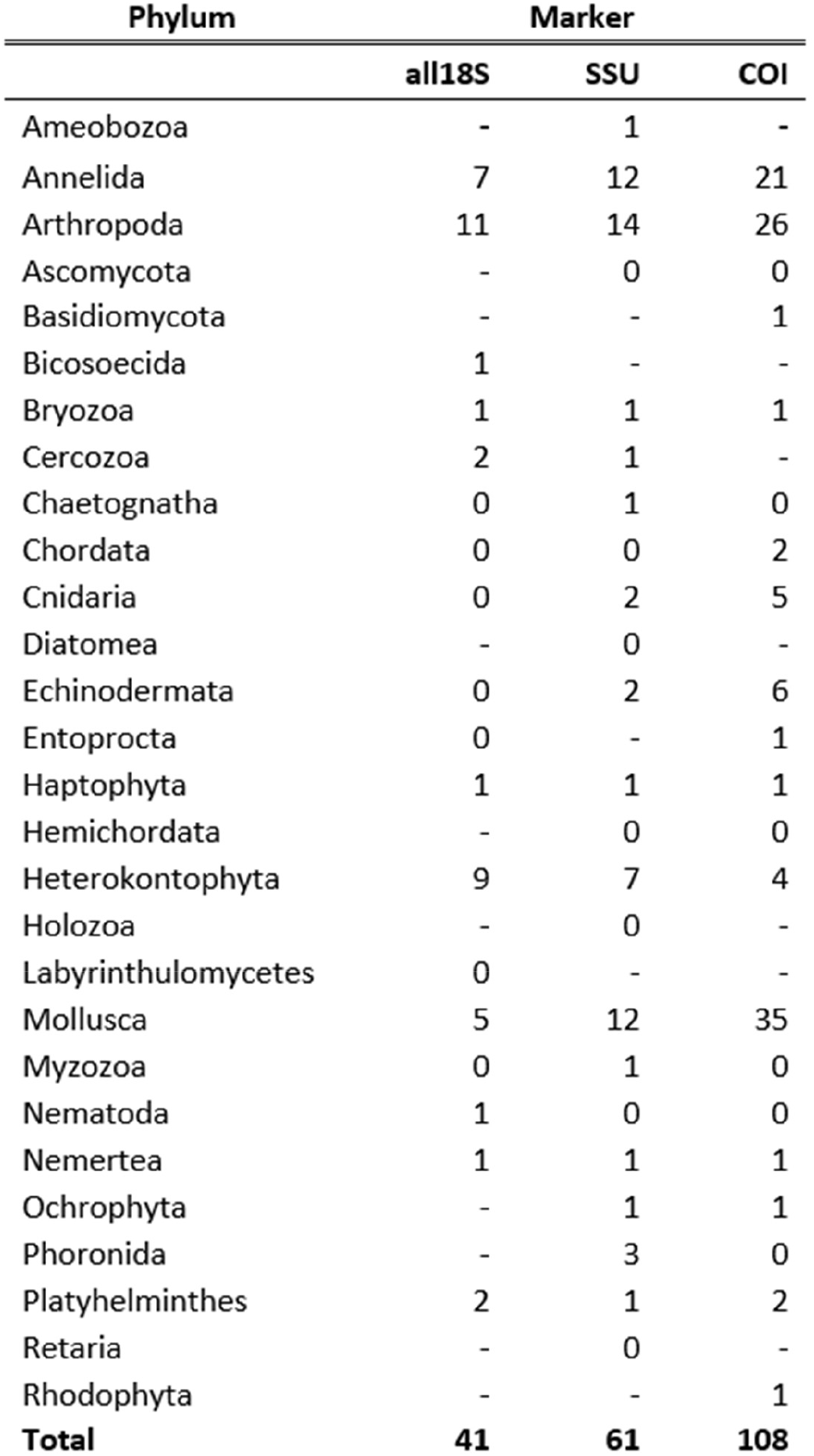
Number of species-level assignments in each phylum per marker. A value of 0 indicates that members of the phylum were present but were not identified to the species level.

The proportion of ASVs that could be given species-level assignments also varied across loci: 17.7% of ASVs could be given species level assignments for all18S, 29.5% for SSU, and 22.1% for COI primers. Most species level identifications were within the Annelida, Arthropoda, and Mollusca phyla (Table 4). Of these species level assignments, very few were identified with all primer sets; for example, only two species were identified with all three markers (Fig. 2d). However, the two 18S primers had higher numbers of species in common than the COI marker did with either of the 18S markers (Fig. 2d).

Notable differences in the relative abundance of each phylum occurred between the three markers (Figs. 3 and 4). For example, in an examination of the top 10 most abundant phyla, the 18S primers detected a higher median relative abundance of Arthropoda (all18S = 79.4%, SSU = 73.6%) than the COI amplicon (30.7%) (Fig. 4). The median relative abundance of Chaetognatha (arrow worms), in comparison, was greatest in the COI dataset (59.5%), while members of the Chaetognatha phylum made up substantially less of the total relative abundance in the all18S and SSU datasets (median values = 11.5% and 17.4%, respectively) (Fig. 4). Similar, but less considerable variations in the relative abundance of the other co-occurring phyla were also evident across the three markers (Figs. 3, 4, and S2). For instance, the Heterokontophyta (golden and brown algae) were far more common in the SSU dataset, comprising between 20% and 30% of some samples. Some phyla were present only in the 18S or COI datasets: members of the protist phylum Cercozoa were only identified by 18S markers, while Basidiomycota (fungi) and Rhodophyta (red algae) were exclusively identified by the COI amplicon. The relative abundance of taxa unassigned at the Kingdom and Phylum levels was remarkably low across all markers, with the greatest median relative abundance of taxa unassigned at these levels equating to only 0.62% of the total relative abundance in the COI dataset (Fig. S4).

**Figure 3.**
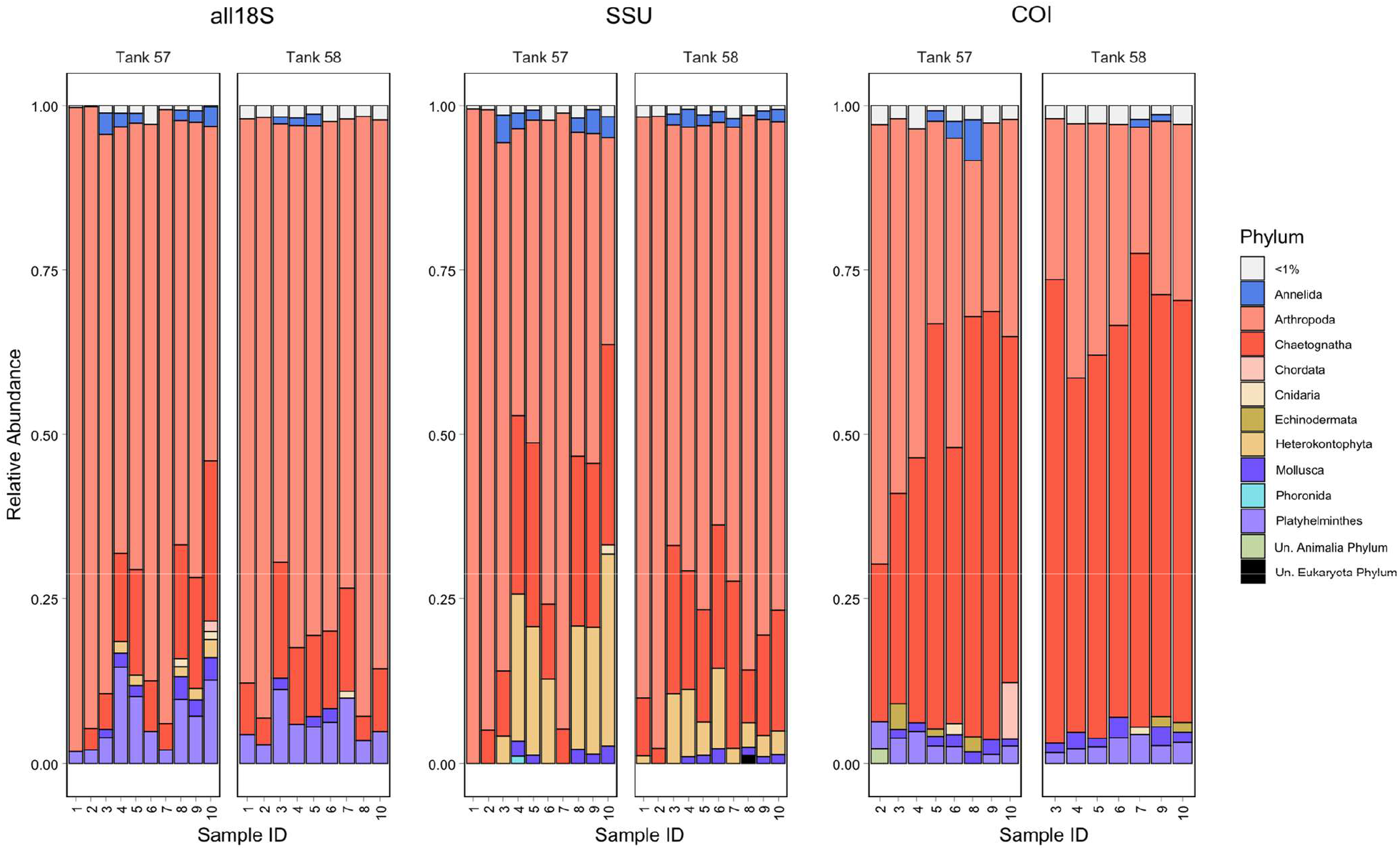
Barplot showing relative abundance of the top 10 phyla across all loci. Each vertical bar represents one sample, samples are clustered by tank and marker as indicated. All other phyla except for the top 10 most abundant are combined as “other.”

**Figure 4.**
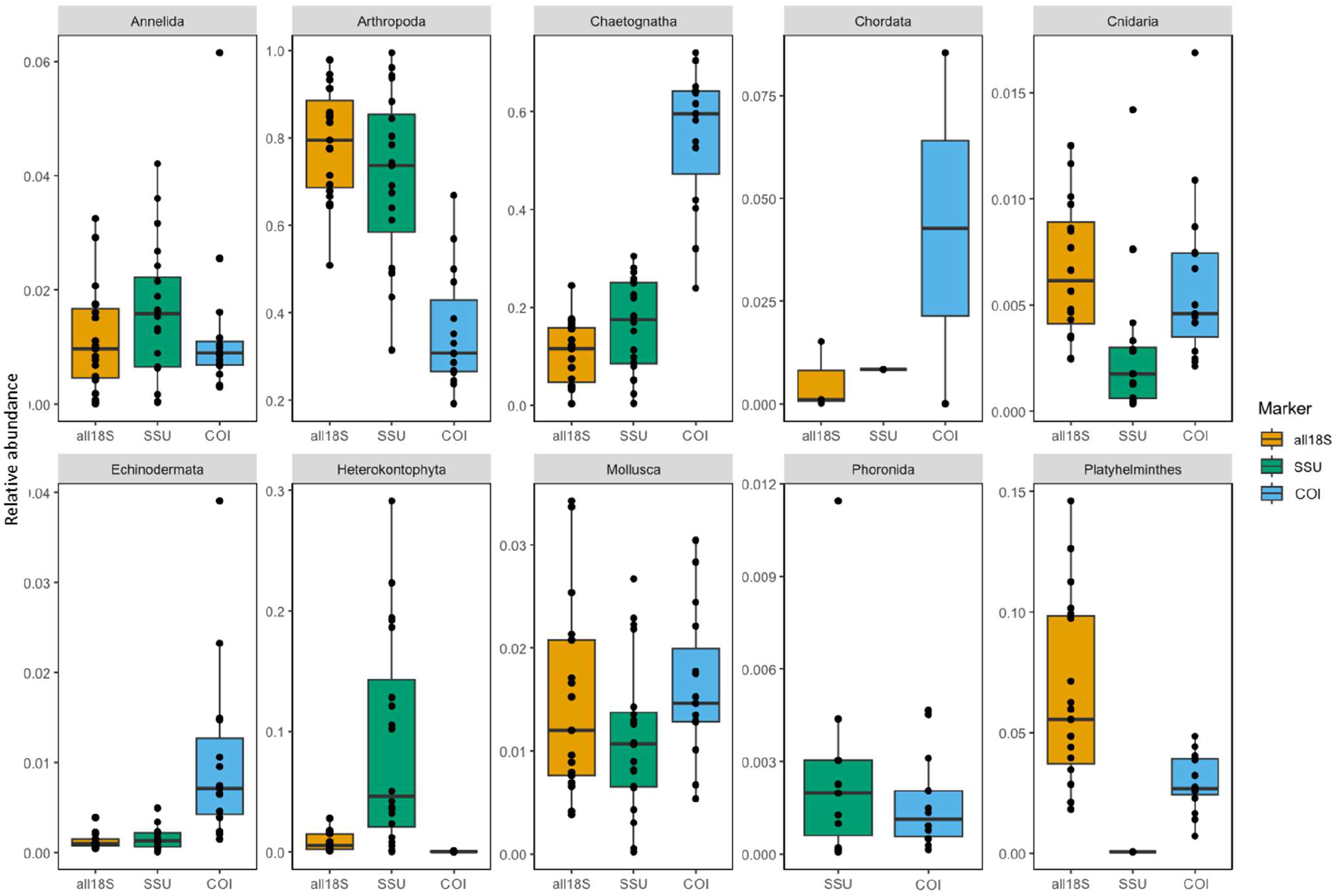
Box plots showing the relative abundance of the top 10 most abundant phyla across all markers, based on data rarefied at 14,156 reads.

Species with a history of anthropogenic introduction were identified within all marker datasets, based on comparisons of marker species lists with the NEMESIS database (Table 5, Table S3). However, the number of reads per species and the confidence level of these assignments varied considerably (Table S3). The COI marker revealed many more non-native taxa than the 18S markers, and included species in the phyla Annelida, Arthropoda, Bryozoa, Chordata, and Mollusca (Table 4). Trends in rarefaction and extrapolation curves of introduced species mirrored those of the full dataset; rarefaction and extrapolation curves approached plateau for the SSU and COI markers, with 99% of the total estimated non-native species biodiversity detected with the SSU marker; however, less than 95% of the estimated non-native species biodiversity was detected for the COI marker (Fig. 5, Table S4).

**Table 5.**
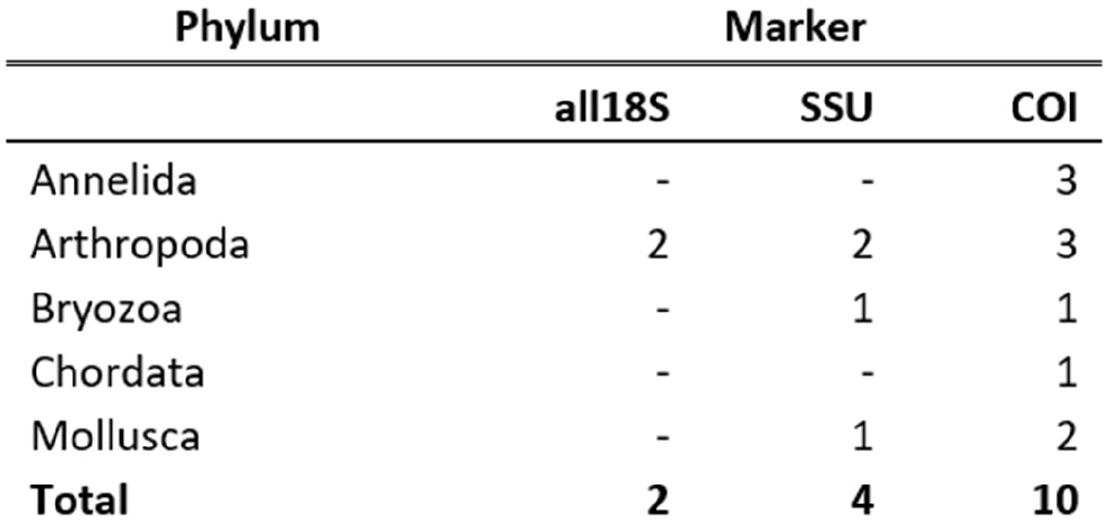
Number of observed non-native species in each phylum per marker. An empty space indicates that no invasive species were identified in that phylum.

**Figure 5.**
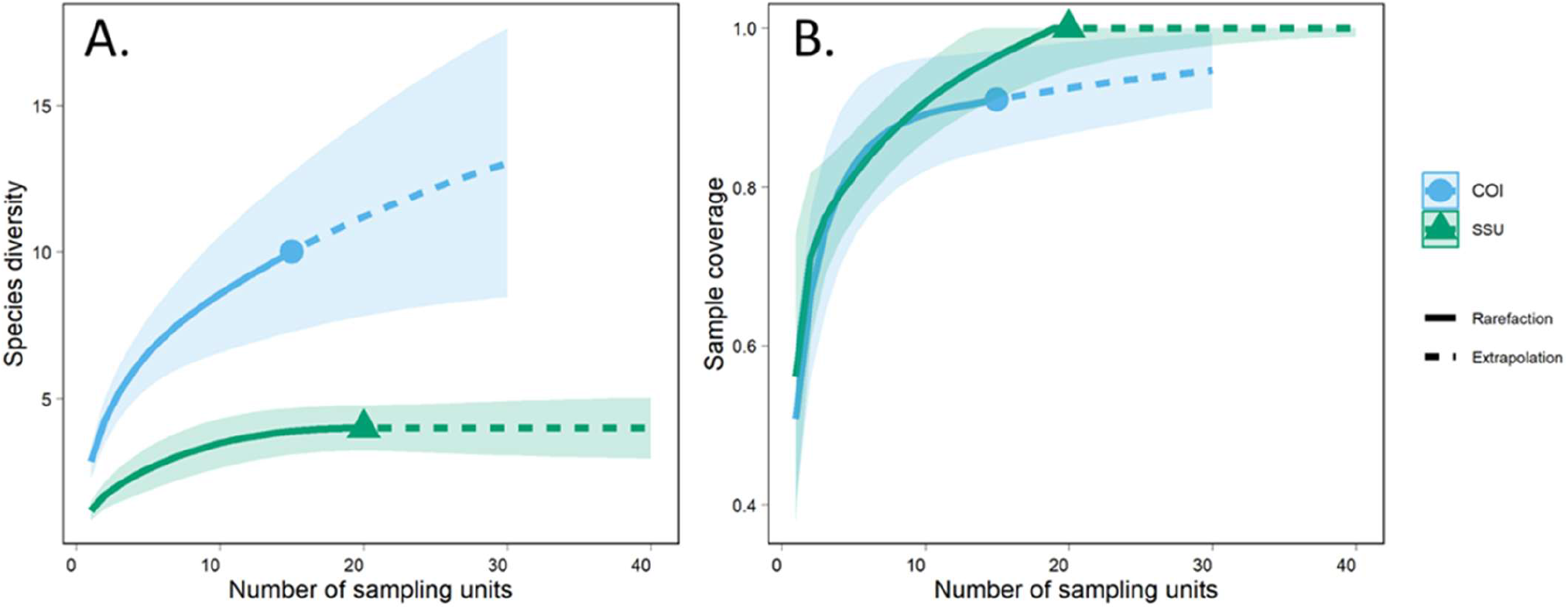
Accumulation curves for non-native diversity based on an iNEXT analysis of data rarefied at 14,156 reads. A. Diversity accumulation curves, with species-level diversity on the y axis and number of sampling units on the x-axis. B. Sample coverage, with the fraction of total estimated species diversity on the y axis and number of sampling units on the x-axis. Data for different loci are plotted separately, as shown in legend. Solid lines indicate interpolation based on observed data, dashed lines represent extrapolated estimates, shaded areas show 95% confidence intervals. The all18S primers could not be interpolated due to a low number of observations. Total estimated non-native species richness was 17 for COI, and 4 for SSU.

## 4. Discussion

By our most conservative estimate there are approximately 272 identifiable species from 28 eukaryotic phyla in the BW of this vessel. This is based on rarefaction and extrapolation of a combined dataset including ASVs from all three primer sets to which we can assign species names. Limitations in reference data preclude us, however, from identifying most ASVs at that level of taxonomic resolution. Given that species-level assignments were available for approximately 25% of ASVs across all markers, a more liberal estimate would place the overall total diversity at upwards of 1000 species. This is consistent with our assessment of overall molecular diversity. Of the over 1700 total estimated OTUs (742 for COI, 544 for all18S, and 425 for SSU), we cannot know what proportion might be shared across multiple markers. However, given that each primer set recovers different taxonomic diversity at both the phylum and species levels (Fig. 2b and 2d), some fraction of this OTU diversity is surely unique to individual markers, and it would not be surprising if total unique OTU richness were well over 1000 eukaryotic taxa.

While these estimates are from only a single ship, we have no reason to believe that this vessel harbors an unusually diverse BW community. The vessel analyzed in this study was sampled after a 7-day voyage from Long Beach, California, without any BW management. Traffic into the port of Valdez generally consists of US flagged ships that carry single-source water from only a few US west coast locations [64]. Such ships are expected to carry lower diversity and are less likely to exhibit significant differences in diversity between tanks than vessels with more complex ballasting histories across diverse ports and global regions. In fact, a previous study found that alpha diversity measures on this and similar vessels entering Valdez were lower than those for vessels entering two other US ports on the Atlantic and Gulf coasts [23].

Our overall richness estimates are substantially higher than suggested by previous assessments of BW diversity. Not surprisingly, they are roughly an order of magnitude higher than the taxonomic richness observed in morphological studies, which have largely been limited to metazoan diversity. Such studies typically have reported on the order of one to two dozen species per vessel [13-15, 17, 65]. Even studies agglomerating across large numbers of vessels have not described comparable levels of diversity; for instance, Cordell et al required 380 ballast tanks (from multiple sources, seasons, and ships) to observe 124 total zooplankton taxa via morphological analysis [16]. Assessments employing multi-locus metabarcoding approaches have generally revealed considerably higher richness estimates, but even in those cases we are not aware of comparable levels of eukaryotic diversity being reported from a single vessel (we ignore here studies that have explored bacterial diversity in BW, e.g. [66-71]). Zaiko et al did utilize a multi-locus approach to evaluate diversity from a single ship, which was sampled multiple times over the course of a 3-week voyage [25]. That study reported 29 OTUs assigned genus-level taxonomy from one locus and 136 from another. Ardura et al conducted a similar investigation exploring changes in diversity throughout the course of a voyage, identifying 165 OTUs across two markers from one vessel and 631 OTUs on another [24]. More common have been assessments of diversity across multiple ships that use a single marker region. In these cases, extremely high levels of cumulative molecular diversity have often been reported across ships [23, 26, 27], but it is difficult or impossible to determine per capita richness.

Importantly, none of these previous investigations had the explicit aim of describing the total diversity present in a single vessel. In the current study we took advantage of an intensive sampling approach and the variation present between samples to estimate total taxonomic richness more thoroughly. Our observation of differences between samples (Fig. 3) is expected given the known patchiness of BW biota and the possibility of stochastic variation introduced by metabarcoding. Like any aquatic habitat, BW tanks provide multiple opportunities for organisms to distribute themselves non-randomly; this may occur through response to physical forces (e.g. depth) or biological interactions (e.g. aggregation) [2, 15, 72]. Variation in community composition from sample to sample is therefore anticipated. In addition, metabarcoding studies can exhibit significant variation among even technical replicates, in large part due to stochastic extraction and amplification of rare taxa [73, 74]. For instance, replicate samples taken from even the same locality can yield slightly different assessments of community composition [75], and as many as 9 PCR replicates could be necessary to get complete estimates of biodiversity using COI metabarcoding [76]. It is precisely this variation that allows for fuller appreciation of the biodiversity in these samples.

The intensive sampling approach enables us to capture a substantially greater fraction of the rare diversity that exists in these tanks. This is additionally facilitated by our ability to distinguish diversity that typically would not be amenable to characterization via traditional methods. The power of metabarcoding approaches to capture such “hidden diversity” is well-known and has dramatically expanded expectations regarding overall levels of biodiversity in various environments [77, 78]. This is particularly true when multiple genomic loci are employed to cast the taxonomic net as widely as possible [37, 79, 80]. Our analyses, for example, capture microbial eukaryotic diversity far more effectively than previous morphological studies, including representatives of single-celled fungal, algal, and protistan phyla such as Oomycota, Ascomycota, Cercozoa, Amoebozoa, Haptophyta, Bacillariophyta, and Dinoflagellata. The value of metabarcoding in accounting for such non-metazoan diversity has been recognized in previous studies [41, 42, 81], and while interest in potentially harmful BW-borne invasions has prompted morphological studies focused on a subset of these taxa (e.g. diatoms and dinoflagellates [82]), broad-based morphological assessments of BW diversity have typically been limited to metazoans.

The adoption of multiple primer sets in our analysis further enables a more comprehensive assessment of total diversity. For taxa identified at the species level, our total estimated richness across all three markers was considerably higher than that estimated for the most diverse single marker alone (272 total vs. 209 for the COI primer alone, Fig. 1C vs. E). The vast majority of species, in fact, were only identified with a single primer set (Fig. 2D), despite substantial overlap in taxonomic coverage at the phylum level (Fig. 2B). Significant differences in taxonomic coverage and biodiversity recovery were even observed between the two primer sets targeting the 18S rRNA locus, consistent with previous observations [83].

Our analyses suggest that the COI primer set is the most effective for describing overall diversity, as it recovers the greatest taxonomic breadth at both the phylum and species level, although we achieved a higher percentage of species-level assignments with the SSU primer set (29.5% vs. 22.1% for COI). This seems in keeping with standard practice and general experience indicating the value of COI for barcoding of metazoans [37, 71, 84]. It is perhaps somewhat surprising that our COI primer set captures just as much taxonomic diversity at the phylum level as our 18S primer sets, since the latter have generally been shown to offer more holistic representation across the tree of life [37, 85]. Nevertheless, for overall biodiversity assessment, our results support recommendations for the application of multiple markers in metabarcoding studies, assuming the availability of resources, particularly when the goal is capturing the total available diversity [33, 71, 86-88].

Our results suggest that the scale of biodiversity transported in BW may have been dramatically underestimated. It has been reported that more than 120,000 vessels are in active service, approximately 52,000 of which would typically carry BW (e.g. tankers, cargo ships, container ships, etc.) [91]. If we conservatively estimate that only 75% of those active vessels is on the water on any given day, we would expect to find 40,000 BW-bearing ships traveling the world’s oceans daily. Our most conservative estimates suggest that 272 identifiable species might be carried in the BW of a single vessel; at the higher end of our estimates, we would not be surprised to find well over 1000 different taxa. Other studies have shown that hundreds of vessels may be required to fully sample even regional biota [23], so while common species may comprise the bulk of the diversity delivered in BW, the constant presence of rare taxa suggests that each vessel is likely to deliver some proportion of unique species. If we assume that proportion to be even as low as 1% (conservatively, 2.7 species), we extrapolate to approximately 108,000 species hitchhiking daily across the world’s oceans in BW; yet, given the global scale of shipping and BW sources across distinct biogeographic regions, this may still underestimate daily flux of BW diversity. This perhaps alarming estimate, if true, invites two critical observations. First, global legal frameworks aimed at reducing the risks that these taxa might survive transit and establish outside their native ranges are critically important. The international consensus on BW management [9], in particular, has been a remarkable achievement and is diminishing the astonishing propagule pressure delivered to recipient coastal waters. Second, it is highly likely that this extraordinary exchange of biodiversity has resulted in countless human-mediated invasions (including range expansions) that have yet to be recognized, with completely unknown consequences [21]. Integrating multi-locus metabarcoding and intensive sampling efforts into future BW studies could therefore enable more efficient detection of previously undiscoverable or rare biodiversity, including non-native species, thus providing more accurate estimates of overall BW diversity. Future focus on previously under-studied components of BW diversity (e.g. microscopic eukaryotes, parasitic taxa, etc.) using molecular methods could identify taxa that have likely been translocated via BW and introduced globally, potentially providing novel insights into the richness and diversity of non-native species impacting coastal systems.

While multi-locus metabarcoding represents a remarkable advance in our ability to assess overall biodiversity, the approach is still burdened by several important limitations. For instance, the inadequacies of available reference data have been widely acknowledged and many taxa are absent from even the most ardently populated databases, preventing accurate assignment of taxonomy for many sequences [31, 51, 94]. This likely explains, in large part, the fact that only a fraction of the diversity observed in our datasets could be assigned species-level taxonomies. Even when reference sequences exist, bioinformatics pipelines are myriad and complex, and errors in assignment can occur [95].

Here, we have focused on overall biodiversity, but still caveats are warranted. For species-level assessments, we are necessarily discarding a large proportion of our data which cannot be confidently assigned at those taxonomic levels. Still, without additional scrutiny, it is difficult to fully assess confidence in the assignments that *are* retained, and some may be erroneous. For OTU-level estimates of diversity we note that, while clustering at 97% identity is meant to reflect typical species-level genetic variation, this may over- or under-represent true diversity depending on taxonomic group [84, 96]. Nevertheless, we believe that the multiple estimates offered in our analyses fairly represent the overall diversity present in these samples. It is also important to recognize that our results specifically capture molecular diversity; while this is reasonably taken to reflect the organismal diversity in our samples, it is impossible to determine how many of the sequences observed actually derive from live organisms [70, 97]. DNA could enter a BW tank through entrainment of dead organisms (or even their shed environmental DNA) or as stomach contents of entrained predators, or organisms present in tanks could die in transit and shed their DNA postmortem. Persistence of nucleic acids in BW tanks has not been well studied, but it is possible that it may be considerable given physical characteristics of the environment [98, 99].

These considerations are particularly important when attempting to extract information on taxa of concern, such as introduced and invasive species, from metabarcoding datasets [100]. Our analyses indicate the likely presence of some non-native taxa in the BW of this vessel (Table 5), and total estimated non-native species richness was as high as 17 species for the COI primer; inclusion of multiple markers increased the number of non-native taxa observed, consistent with previously published studies [101, 102]. However, since we are screening only against non-native taxa known to be present in the continental United States (from the NEMESIS database), this estimate of total non-native diversity is by at least one measure quite conservative. Even within the small fraction of diversity that can be identified to species level, there are likely additional taxa that are non-native to the recipient region and yet not captured as such in the NEMESIS database. And we are obviously disregarding any molecular evidence of non-native taxa for which there is no existing reference data to allow taxonomic assignment [103]. For those non-native taxa that we have identified, additional bioinformatics assessment would be necessary to better evaluate confidence in those taxonomic assignments [104]; even then, our analyses could only provide molecular evidence of those species in the samples and could not demonstrate the presence of live individuals. The analyses presented here thus offer only a very limited assessment of the potential direct risk of non-native species introduction posed by this and similar vessels.

While our estimates suggest that BW-borne diversity may be much higher than previously thought, it is also important to recognize that is likely a substantial *under*estimate of the total diversity carried by this vessel. First, we sampled a relatively limited portion of the overall eukaryotic biodiversity present on a ship. For example, since we used 80 μm zooplankton nets to collect these samples, we are likely underestimating the diversity of microbial eukaryotes and are only including those associated with zooplankton taxa. We have also not explored (a) BW sediments, which may contain considerable diversity and differ from that present in the water column [105] or (b) biofouling organisms present on the hull and underwater surfaces of the ship; in particular, biofouling may be just as potent a vector of species introduction as BW [22], and assessments of taxonomic richness on ships’ hulls suggest that this diversity may be substantial and underexplored [106, 107]. Second, while we adopted an intensive sampling approach in order to maximize detections of biodiversity, we ultimately did not achieve saturation of the species accumulation curves (Fig. 1, Table 2), indicating that additional sampling should reveal additional biodiversity. Finally, given differences in the biodiversity of BW originating from different regions [23, 26, 41] and the lack of comparable intensive sampling studies, we cannot assume that our estimates of BW diversity in a single vessel are representative of all BW. Rather, our results serve as an example of the possible diversity present in a single vessel and indicate the value of repeated sampling and multi-locus metabarcoding for biodiversity assessments in BW. Noteworthy gaps thus exist regarding the full potential of vessels as vectors [108], and while our study advances our understanding of vessel-borne biodiversity it nevertheless remains an incomplete picture.

## 5. Conclusions

This study presents the first reported use of a combined intensive sampling and multi-locus metabarcoding approach applied to the BW of a single vessel. Our analyses revealed the presence of 272 identifiable species from 28 eukaryotic phyla, with estimates of the overall total diversity reaching upwards of 1000 species. Although our results provide much higher estimates than previous studies of the total eukaryotic diversity being transported in BW, it is likely that we are still underestimating BW biodiversity. These results not only provide a novel and revealing insight into the scale of biodiversity being transported globally in BW, but also further demonstrate the utility of multi-locus DNA metabarcoding as a tool for characterizing that biodiversity and for assessing risks associated with BW and other vectors of anthropogenic introductions.

## Supplementary Materials

see associated file “Supplementary Tables & Figures.”

## Author Contributions

All authors contributed to conceptualization and planning of the work described. JKC and KKH led sampling efforts. SB, EMP, and JAD generated and curated the data. SB conducted the formal analyses. SB and JAD wrote the initial draft. All authors reviewed and edited the final draft. All authors have read and agreed to the published version of the manuscript.

## Acknowledgments

This work was supported by funding from US Coast Guard. The authors wish to thank Alyeska Oil Terminal and their staff and vessel crew for providing access and support to sample ships’ BW, and Mara Cuebas Irizarry and Eric Villegas for invaluable comments on an earlier manuscript draft. The views expressed in this article are those of the authors and do not necessarily reflect the views or policies of the U.S. EPA.

