## Supplemental figures and tables for "Multi-locus metabarcoding and intensive sampling reveal extraordinary diversity carried in the ballast water of a single vessel"

### Supplemental Tables and Figures

**Table S1**. Results of Wilcoxon Rank Sum Tests examining differences in Shannon diversity and OTU richness (Observed) between the two tanks for each amplicon.

| | | W | p-value |
|---|---|---|---|
| **all18S** | Shannon | 61 | 0.211 |
| | Observed | 50 | 0.7129 |
| **SSU** | Shannon | 70 | 0.1431 |
| | Observed | 63.5 | 0.3252 |
| **COI** | Shannon | 42 | 0.1206 |
| | Observed | 47.5 | 0.02706 |

**Table S2**. P-values from pairwise Wilcoxon Rank Sum tests comparing a) OTU richness, and b) Shannon diversity between amplicons. Asterisks indicate significant p-values.

a.

| | **all18S** | **SSU** |
|---|---|---|
| **SSU** | 0.23200 | NA |
| **COI** | 0.000001* | 0.0000012* |

b.

| | **all18S** | **SSU** |
|---|---|---|
| **SSU** | 0.0327* | NA |
| **COI** | 0.0001271* | 0.00* |

**Table S3.** Non-native species identified in each marker dataset, with the confidence level of the species level taxonomic assignment and the total number of reads for each species. The Feature ID is the identifier associated with each ASV in the QIIME2 metadata.

| | Feature ID | Phylum | Species | Confidence level | Total number of reads |
|---|---|---|---|---|---|
| | 213578c85d82b2f9c79ed8669a4282ee | Arthropoda | Ciona intestinalis | 0.8055 | 24 |
| | b179b0520c16365351a6ff90fee325ef | Mollusca | Argopecten irradians | 0.7598 | 12 |
| | 0c1d1f50c103dd7481e016fede839aa3 | Bryozoa | Anguinella palmata | 0.9997 | 7 |
| | 506a7a326aa5e8db8214a47002ccde84 | Bryozoa | Anguinella palmata | 1.0000 | 4 |
| **SSU** | 40f934c5d033632e89ea9af747f4f953 | Arthropoda | Pseudodiaptomus marinus | 0.9910 | 61 |
| | 4a2f850526e9c4c4bcb28555c6bde4e6 | Arthropoda | Pseudodiaptomus marinus | 0.9909 | 542 |
| | 6cc24d6367f6d9486fdee9ec6743d0f6 | Arthropoda | Pseudodiaptomus marinus | 0.9578 | 1054 |
| | d3bf937afe029c26fac9ab6b982d3b11 | Arthropoda | Pseudodiaptomus marinus | 0.8714 | 182 |
| | 4cc20d275239115ace3d2cb1051774e7 | Arthropoda | Pseudodiaptomus marinus | 0.7878 | 2 |
| | 38a06d840bfcbac279d356fdff027ee5 | Arthropoda | Amphibalanus amphitrite | 0.8913 | 3 |
| | f3f9979a4cf7c7900d8822c1e5958dc1 | Arthropoda | Pseudodiaptomus marinus | 0.9992 | 140 |
| | 9e91f29a96632e07fd12c696465f7bf8 | Arthropoda | Pseudodiaptomus marinus | 0.9992 | 119 |
| | aece0f5b019fbb9e53e3d0c458f4047f | Arthropoda | Pseudodiaptomus marinus | 0.9992 | 58 |
| | 0180b2135d01de964e5691b79cb3dafb | Arthropoda | Pseudodiaptomus marinus | 0.9993 | 64 |
| | 8e385c77f19c659b6616b2aa2b78b80b | Arthropoda | Pseudodiaptomus marinus | 0.9951 | 29 |
| | 939632f0297b2ecb7b494c93a6c0c15a | Arthropoda | Pseudodiaptomus marinus | 0.9951 | 8 |
| | f887af34f5cb52d9fa5a7f6c3aa25fc1 | Arthropoda | Pseudodiaptomus marinus | 0.9951 | 65 |
| | 2979f27477eb633bf37d112fa88e1208 | Arthropoda | Pseudodiaptomus marinus | 0.9946 | 43 |
| | a91356030345b9ec76f5e7cfb70b274d | Arthropoda | Pseudodiaptomus marinus | 0.9947 | 41 |
| | 2e457db0b38f1fed24c32e157a735d81 | Arthropoda | Pseudodiaptomus marinus | 0.9949 | 40 |
| | 29dcc432ddf53f81f37de3cfe6c7e8cf | Arthropoda | Pseudodiaptomus marinus | 0.9785 | 36 |
| | a9bf7838eb06da648280f22a68d47be5 | Arthropoda | Pseudodiaptomus marinus | 0.9993 | 60 |
| | 3ccc553f6b0d67cc064f02162e33f83a | Arthropoda | Pseudodiaptomus marinus | 0.9993 | 27 |
| | cf283decc12ea8e6124ea6eaed05fbb4 | Arthropoda | Pseudodiaptomus marinus | 0.9993 | 17 |
| | 39566213331888538ddc5459d613ac9d | Arthropoda | Pseudodiaptomus marinus | 0.9992 | 13 |
| | 335a57eead236ad08abc6a44a1ddf880 | Arthropoda | Pseudodiaptomus marinus | 0.9978 | 36 |
| | fb544fb32abbe5a5e24512e259a37b4c | Arthropoda | Pseudodiaptomus marinus | 0.9979 | 30 |
| **all18S** | 9ca1f7ff55ad68b001195a483308b341 | Arthropoda | Pseudodiaptomus marinus | 0.9996 | 43 |
| | 0652e2623724bc806b6f743103992cb3 | Arthropoda | Pseudodiaptomus marinus | 0.9996 | 73 |
| | f73965f5bce9b7260e29537bd1b010d1 | Arthropoda | Pseudodiaptomus marinus | 0.9993 | 34 |
| | e44092ebf1f0d1a4ed90f0fc90d12136 | Arthropoda | Pseudodiaptomus marinus | 0.9994 | 27 |
| | c573c0d0c540b19f42e4b19aeb73a752 | Arthropoda | Pseudodiaptomus marinus | 0.9994 | 28 |
| | 87c3c333e402a85dd54d108e1bf60c48 | Arthropoda | Pseudodiaptomus marinus | 0.9994 | 9 |
| | 9e865ba5700a5defff8a860c01ebed17 | Arthropoda | Pseudodiaptomus marinus | 0.9994 | 53 |
| | 4546db6d3ecf316f050397023a9e2277 | Arthropoda | Pseudodiaptomus marinus | 0.9994 | 33 |
| | eeac049acaffaedbb0d8ee1b6325dca7 | Arthropoda | Pseudodiaptomus marinus | 0.9994 | 47 |
| | f19b2acd3136f993f45374553d0f3b10 | Arthropoda | Pseudodiaptomus marinus | 0.9994 | 25 |
| | 565e81d4d38da8a44c3fa47dd7941a62 | Arthropoda | Pseudodiaptomus marinus | 0.9993 | 19 |
| | 2b33a3a7845f38169e2bc8d70031907d | Arthropoda | Pseudodiaptomus marinus | 0.9965 | 8 |
| | e34e4ca60b439356157d3cd121337c5f | Arthropoda | Pseudodiaptomus marinus | 0.9962 | 21 |
| | cfb2af6aba7dc9d07c4c3fe090a028cc | Arthropoda | Pseudodiaptomus marinus | 0.9964 | 41 |
| | f4d783161ccddc511cd4b226fbdce65d | Arthropoda | Pseudodiaptomus marinus | 0.9960 | 24 |
| | 63c608a831c64b5279e607cdfeee09c4 | Arthropoda | Pseudodiaptomus marinus | 0.9960 | 10 |
| | 1e50d92d032c2d145dae37b8773f64b0 | Arthropoda | Pseudodiaptomus marinus | 0.9843 | 15 |
| | b3c1abd4aee19c7d207bf87bf38dc8b0 | Arthropoda | Pseudodiaptomus marinus | 0.9849 | 8 |
| | a4393809a9476fc5282976dcb9a34cec | Arthropoda | Pseudodiaptomus marinus | 0.9849 | 36 |
| | 69f146e25359278c21051711e948ce58 | Annelida | Hydroides elegans | 1.0000 | 7 |
| | 98fc6233e9aee07b3d1d86808be510b0 | Mollusca | Theora lubrica | 1.0000 | 39 |
| | 6d7d0457fe7c46cc1d2ba20d5e6c4403 | Mollusca | Theora lubrica | 1.0000 | 3 |
| | 5e2b8ef539876d9c37bdfdbf797959f2 | Mollusca | Philine auriformis | 1.0000 | 8 |
| | 1748c13dbef4be5397d86ac3cb58bb91 | Mollusca | Philine auriformis | 1.0000 | 5 |
| | 0088fd1659cbed03babedf420ce8fcca | Mollusca | Philine auriformis | 0.9990 | 8 |
| | 93968353e78cbde33ef1542359c07d26 | Mollusca | Philine auriformis | 0.9985 | 2 |
| | 359e6ca0da39a64e0e205dc2b0aadc9e | Bryozoa | Membranipora membranacea | 0.9451 | 16 |
| | 37a0a4b86e4fc2e8883e403c37272f4a | Arthropoda | Amphibalanus amphitrite | 1.0000 | 3 |
| | 814aa35f82a0c7044f0b4e1d68d21bdd | Chordata | Dorosoma cepedianum | 0.9401 | 1 |
| **COI** | bcf1bf2e180c1d5d6076bcb96115c3d5 | Annelida | Myrianida pentadentata | 1.0000 | 818 |
| | a653ebdfff2abe254a4604bd5f6bfe3e | Annelida | Myrianida pentadentata | 0.9986 | 41 |
| | 047a88da022200bb823c845961dcc600 | Annelida | Pseudopolydora paucibranchiata | 1.0000 | 2 |
| | c53143a88d02fe4cb7ded96615189f96 | Arthropoda | Pseudodiaptomus marinus | 1.0000 | 206 |
| | 0d2aafdc27a44feb4e88f2b6ff6b83fb | Arthropoda | Pseudodiaptomus marinus | 1.0000 | 1461 |
| | 9c53f25d7154a3bfeab2f24c67c265de | Arthropoda | Pseudodiaptomus marinus | 1.0000 | 155 |
| | edf230d3cfdfe5fdfe08398b323e331e | Arthropoda | Pseudodiaptomus marinus | 1.0000 | 392 |
| | d66498164ed69f576ae1c39a669ea895 | Arthropoda | Pseudodiaptomus marinus | 1.0000 | 41 |
| | 544c270523ae14923f645a750c8dfab9 | Arthropoda | Pseudodiaptomus marinus | 1.0000 | 3 |
| | a7d1f80825fdce903975113abdef1d3d | Arthropoda | Oithona davisae | 1.0000 | 8 |
| | cc8f7393e761b3bea6269d79e358e085 | Arthropoda | Oithona davisae | 1.0000 | 9 |

**Table S4**. a. Number of samples required to achieve 50%, 90%, 95%, and 99% coverage of invasive estimated diversity for each locus. Asterisks indicate estimates with high uncertainty (estimated number of samples more than twice the observed number). Number of samples rounded to nearest whole number. b. Percent of invasive estimated diversity captured by sample sizes n = 1 through n = 5. Percentages are rounded to nearest whole number.

a.

| Locus | Per cent coverage | | | |
|---|---|---|---|---|
| | **50** | **90** | **95** | **99** |
| **SSU** | 1 | 10 | 14 | 18 |
| **COI** | 1 | 12 | 32* | 76* |

b.

| Locus | Number of sampling units | | | | |
|---|---|---|---|---|---|
| | **1** | **2** | **3** | **4** | **5** |
| **SSU** | 56 | 71 | 76 | 79 | 81 |
| **COI** | 51 | 66 | 74 | 79 | 83 |

**Figure S1**. Differences between BW tanks. a. Species diversity and sample coverage curves for individual BW tanks. As in Figure 1, solid lines indicate interpolation based on observed data, dashed lines represent extrapolated estimates, shaded areas show 95% confidence intervals. b. Species-level NMDS plot based on the Bray-Curtis ordination showing tank differences for each marker, based on rarefied data. Ellipses indicate 95% confidence intervals, asterisks indicate significance of PERMANOVA results.

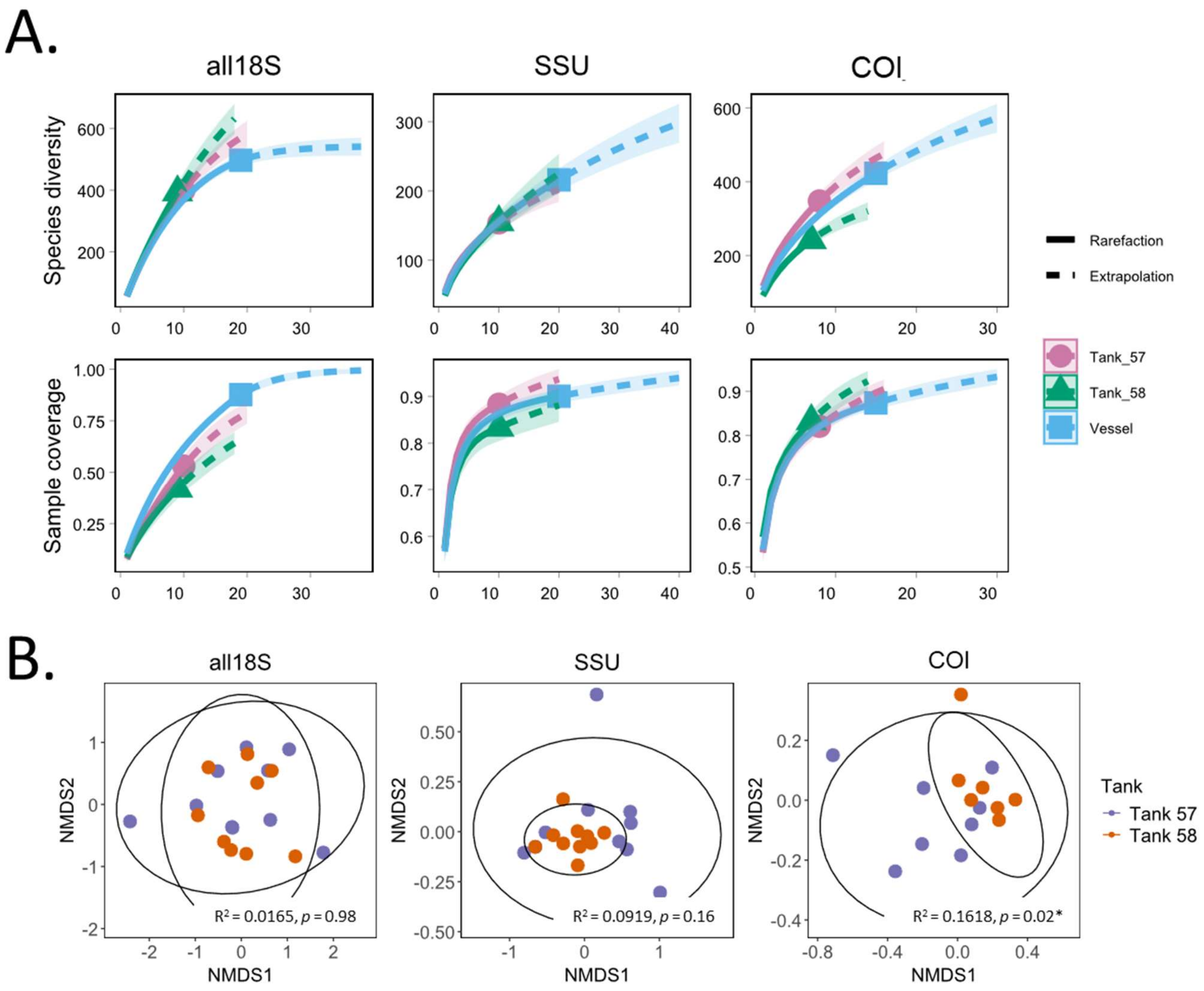

**Figure S2**. A. Average number of reads per marker, after filtering to remove samples not included in the analysis, prior to rarefying. B. Average richness per marker and tank after rarefying, C. Average Shannon diversity per marker and tank after rarefying. Note that the results displayed here are not significantly different between tanks, as determined by a Wilcoxon Rank Sum Test.

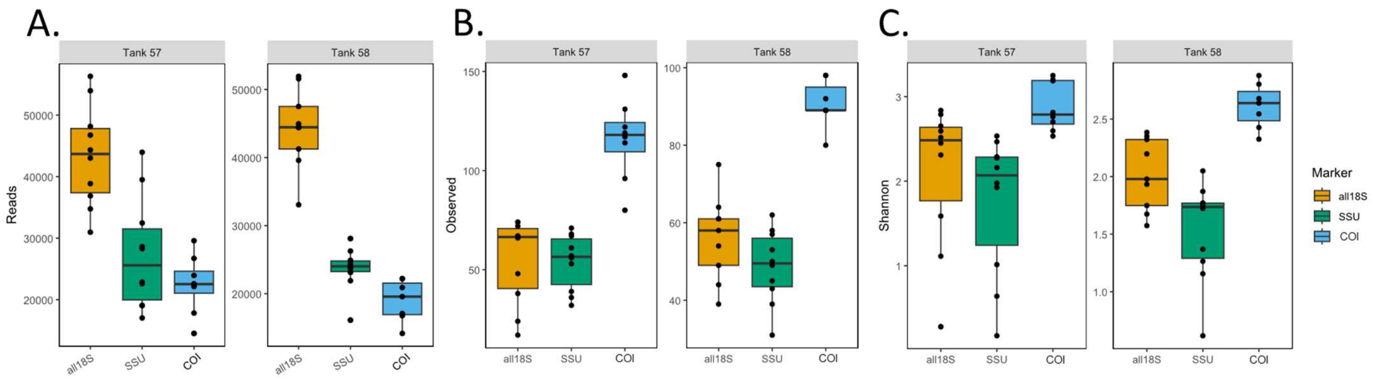

**Figure S3**. A. Average number of reads per marker, after filtering to remove samples not included in the analysis, prior to rarefying. B. Average richness per marker after rarefying, C. Average Shannon diversity per marker after rarefying.

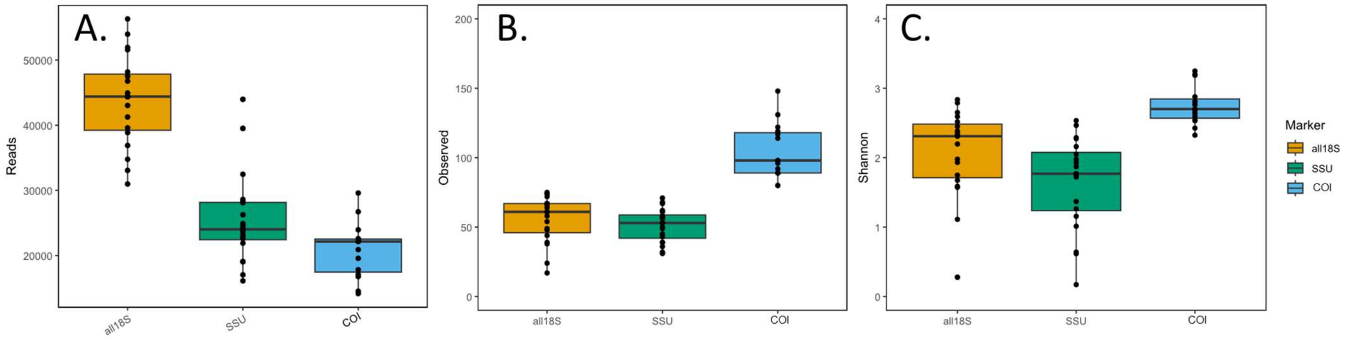

**Figure S4**. Box plots showing the relative abundance of all phyla across all 4 genomic loci, based on data rarefied at 14,156 reads.

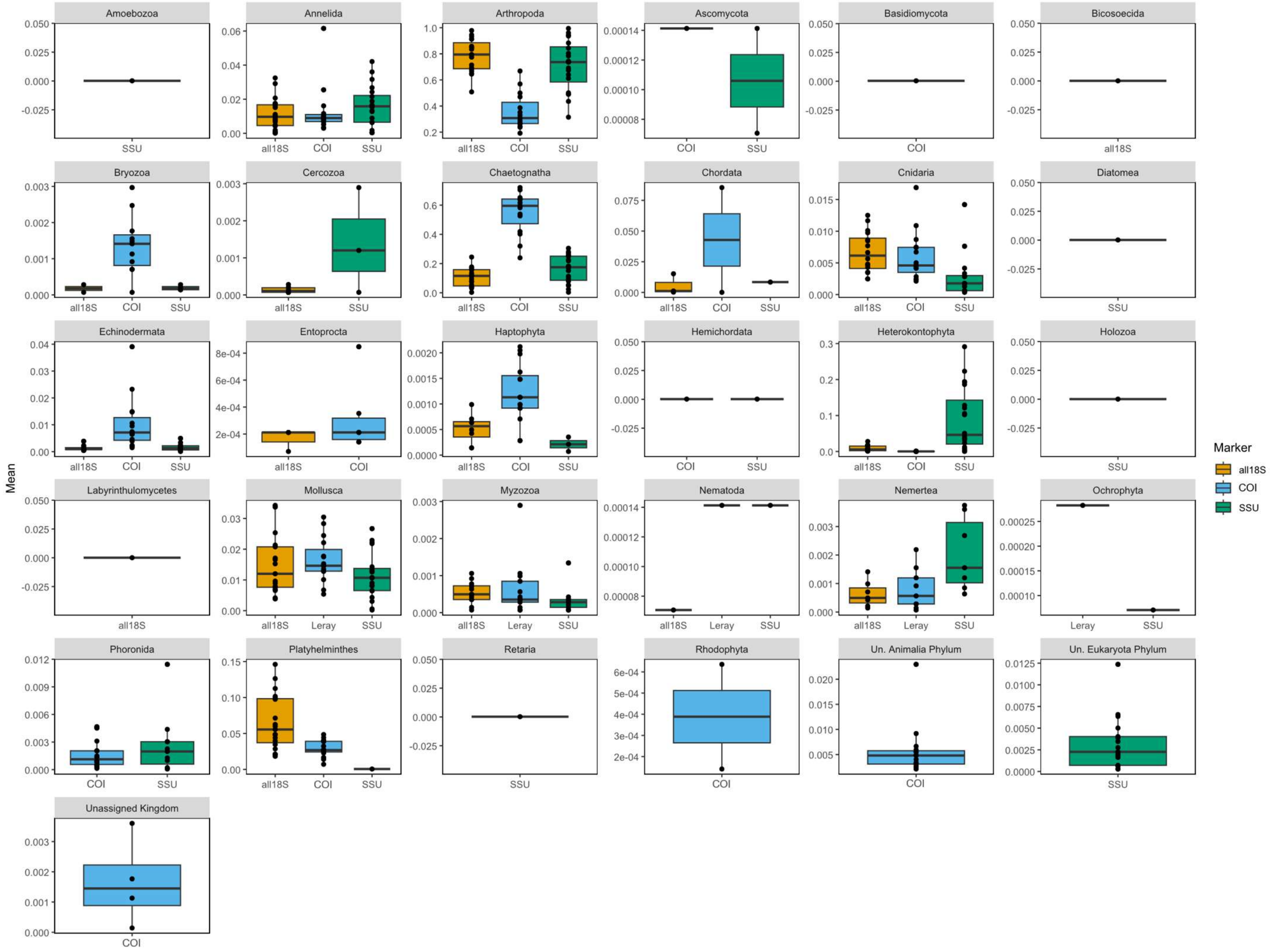
